# Intervene or wait? Modelling the timing of intervention in conservation conflicts adaptive management under uncertainty

**DOI:** 10.1101/2021.08.27.457773

**Authors:** Adrian Bach, Jeroen Minderman, Nils Bunnefeld, Aileen Mill, Alexander B. Duthie

**Affiliations:** Biological and Environmental Sciences department, The University of Stirling, Scotland, United Kingdom; School of Natural and Environment Sciences, Newcastle University, England, United Kingdom

**Keywords:** Adaptive management, Conservation conflicts, Decision-making modelling, Individual-based modelling, Management Strategy Evaluation, Timing of intervention, Uncertainty

## Abstract

The timing of biodiversity managers’ interventions can be critical to the success of conservation, especially in situations of conflict between conservation objectives and human livelihood, i.e., conservation conflicts. Given the uncertainty associated with complex social-ecological systems and the potentially irreversible consequences of delayed action for biodiversity and livelihoods, managers tend to simply intervene as soon as possible by precaution. However, refraining from intervening when the situation allows it can be beneficial, notably by saving critical management resources. Here, we introduce a strategy for managers to decide, based on monitoring, whether intervention is required or if waiting is possible. This study evaluates the performance of this waiting strategy compared to a strategy of unconditional intervention at every opportunity. We built an individual-based model of conservation conflict between a manager aiming to conserve an animal population and farmers aiming to maximize yield by protecting their crop from wildlife damage. We then simulated a budget-constrained adaptive management over time applying each strategy, while accounting for uncertainty around population dynamics and around decision-making of managers and farmers. Our results showed that when the decision for the manager to intervene was based on a prediction of population size trajectory, the waiting strategy performed at least as well as unconditional intervention while also allowing managers to save resources by avoiding unnecessary interventions. Under difficult budgetary constraints on managers, this waiting strategy ensured as high yields as unconditional intervention while significantly improving conservation outcomes by compensating managers’ lack of resources with the benefits accrued over waiting periods. This suggests that waiting strategies are worth considering in conservation conflicts, as they can facilitate equitable management with a more efficient use of management resources, which are often limiting in biodiversity conservation.

## INTRODUCTION

With a growing human population and rising standards of living, the amount of earth’s surface used for human activities is increasingly large and often overlaps with the ranges of species of conservation concern. A conservation conflict can arise when such a species is strictly protected but also impacts human livelihood, potentially leading to a clash of interests over management decisions (Redpath et al. 2013, 2015). Diverging objectives can lead land-users to defect from policies by ignoring or subverting them and engage in illegal activities often hindering conservation objectives (Bainbridge 2017, Bunnefeld et al. 2017, Glynatsi et al. 2018, Rakotonarivo et al. 2020). These conflicts are especially serious when conservation and protection interferes with essential livelihood activities such as agriculture (Behr et al. 2017, Mason et al. 2017). Conservation policies must therefore be in line with land-users’ interests to ensure compliance and maximize conservation success while minimizing the impact on food security and/or farmers’ income. Moreover, because conservation conflicts form complex systems with multiple biological, environmental, geographical, and social components, the response to change in these interlinked social-ecological systems is difficult to anticipate (Wilgen and Biggs 2011, Game et al 2013, Mason et al. 2018). To avoid unforeseen perturbations that might jeopardize biodiversity conservation or human livelihood, management should also embrace the uncertainty around ecological processes and human behavior (Fryxell et al. 2010, Schlüter et al. 2011, Bunnefeld et al. 2011, Cusack et al. 2020).

A practical way to deal with uncertainty challenges and complex systems is adaptive management, a technique seeking to improve management iteratively by learning from its outcomes (Williams et al. 1996, Hicks et al. 2009, Keith et al. 2011). It is particularly well adapted to conservation conflicts management because regular monitoring and policy updates enhance the ability to trade-off between opposing interests (Redpath et al. 2013, Wam et al. 2016, Mason et al. 2018, Richardson et al. 2020). Adaptive management thus tailors the conservation policy as closely as possible to the system’s variations, but when and why to update the policy can be key to better management of social-ecological systems and conservation conflicts (Pérez et al. 2019). Because the consequences of mismanagement can be detrimental and even sometimes irreversible (e.g., crop losses and/or animal population extinction, Kaswamila et al. 2007), conventional wisdom might suggest that conservation success will be maximized by reacting as often as possible with updated policy. But waiting can ultimately lead to better management results when well-planned, because it can bring a variety of benefits, including enhancing knowledge through monitoring or research (Walters 1986, Gregory et al. 2006, Nicol et al. 2018). For example, Sims and Finnoff (2013) modelled the progression of the slow and predicable spread of an invasive species and showed that, due to the knowledge acquired during the period of waiting, a delayed time of first intervention was more efficient in reducing both the spread and damages on the focal ecosystem than intervention immediately after detection of the invasion. In contrast, a delayed intervention when the invasion was fast and erratic caused a loss of control over the species progression, eventually leading to a state where any intervention became pointless. In an adaptive management context, Iacona et al. (2017) modelled national parks’ bird diversity protection schemes and showed that waiting and saving conservation funds to accrue interest before spending it progressively on protection achieved a higher number of protected species and a quicker recovery of the extinction debt than front-load spending. Because financial and human resources for management are often limited (Hughey et al. 2003, McDonald et al. 2008), intervening when the benefits of waiting outweigh the risks can be unnecessary spending, if constraints on conservation funding allocation allows it (Ruiz-Miranda et al 2020, Wu et al. 2020). This trade-off between instances of intervention and waiting in an adaptive management process has not yet, to our knowledge, been explored in the context of conservation conflicts. We hypothesize that by refraining from intervening when conflicting stakeholder interests are already aligned, managers could save resources and use them to enhance impact when intervention will be most needed to deliver conservation and/or land-users’ objectives. We predict that it is likely to be especially relevant in situations where a manager’s lack of resources could be compensated for by benefits accumulated over a period of waiting.

To investigate the effect of the timing of intervention on management quality while accounting for the different sources of uncertainty associated with conservation conflicts, we used the generalized management strategy evaluation framework (GMSE, Duthie et al. 2018). GMSE builds on the management strategy evaluation (MSE) framework, which aims to explore the possible outcomes of alternative management scenarios in order to assess their adequacy to managers’ objectives (Smith 1999). MSE, first developed in fisheries and later for terrestrial species, decomposes the process of natural resources adaptive management over time with sub-models of population dynamics, monitoring, management decision-making and harvesting activities, which inform and influence each other. This structure helps to isolate different components of uncertainty associated with each process when evaluating a scenario (Bunnefeld et al. 2011). GMSE uses an individual-based approach for all four sub-models, simulating uncertainty intrinsically (Grimm 1999, DeAngelis and Grimm 2014), and includes a decision-making sub-model for manager and farmer agents that simulates goal-oriented behavior with the possibility of sub-optimal choices. Furthermore, by generating differences between agents, individual-based models (IBMs) can model another potential source of conflict: the inequitable distribution of costs and benefits among stakeholders. Rakotonarivo et al. (2020, 2021) showed that a higher perceived equity in conservation measures among farmers increased the propensity to choose pro-conservation options. Among-user equity is thus important to model and monitor during conservation conflicts management. Knowing this, we further develop and apply GMSE to evaluate the efficiency of alternative management timing strategies against unconditional intervention and determine whether and how a profitable timing trade-off can be found for conservation conflict management under uncertainty.

We modelled a budget-constrained adaptive management of a conservation conflict in which a wildlife animal population of conservation concern negatively impacts agricultural activities, and farmers can respond by culling to minimize yield loss. We propose two novel timing strategies for the manager to determine whether the situation warrants intervention when the resources saved by waiting generate long term benefits. Through simulations with GMSE, we assessed how each timing strategy affected the quality of management regarding the conflict between biodiversity conservation and agricultural production objectives. We thereby determined for which conditions our alternative strategies resulted in better management than intervening at every opportunity.

## MODEL AND METHODS

### Model overview

#### Model case

To simulate conservation conflict management over time, we develop an individual-based model with a population of wildlife animals (referred to as ‘population’), farmers, and a manager all interacting on an agricultural landscape. The landscape is divided into discrete cells, each of which produces an agricultural yield and can hold any number of animals. Each farmer owns a contiguous block of cells that forms their ‘land’, and the sum of its cells’ productivity determines the farmer’s yield. Animals consume agricultural resources from landscape cells to survive and reproduce, which consequently reduces the farmers’ yield. Farmers can cull animals that are on their own land to reduce yield loss. The manager attempts to avoid extinction by maintaining the population around a predefined target size (*T_N_*), as previously done in, e.g., the management of conflict between mountain nyala antelope conservation and trophy hunting in Ethiopia (Bunnefeld et al. 2013), or between farming and migrating birds’ protection in Scotland or Sweden (Bainbridge et al. 2017, Mason et al. 2017, Nilsson et al. 2021). This target was chosen to be high enough to prevent extinction, but low enough to ensure a satisfactory yield to farmers. The manager’s method is to implement a policy incentivizing or disincentivizing culling as appropriate to get the population size closer to *T_N_*. Hence, following an adaptive management process, the manager updates this policy according to the monitoring of the population size (*N_t_*) at each time step *t*.

#### Manager’s policy-making

The manager receives a fixed, non-cumulative budget *B_M_* at the beginning of each time step, which we interpret to reflect the total time, energy or money available to the manager to implement a change of policy and enforce culling restrictions. The policy is modelled as a cost that farmers must pay to cull an animal on their land. The manager can draw into *B_M_* to raise this cost to discourage farmers from culling and favor population growth and can decrease it to facilitate culling and favor a population decrease. To model the budget needed to enforce a restricting policy, every increase of 1 in the culling cost requires an investment of 10 *b.u* from the manager. Conversely, as the manager does not need to incentivize farmers to remove animals when the policy allows high culling rates, they do not need to spend budget to decrease the cost. The amount by which the manager changes the culling cost is computed by GMSE’s evolutionary algorithm according to their goal that was modelled as minimizing the distance between *N_t_* and *T_N_*.

#### Timing strategies

We explored three timing strategies that determine whether a manager intervenes and updates the policy or waits and leaves it as is. The Control strategy (CTL) was the null model in this study. It corresponds to unconditional intervention at every opportunity and was modelled as the manager updating the policy at every time step. With the Adaptive Timing of Intervention strategy (ATI), we define a permissive range *P_T_* around *T_N_* in the form of *T_N_* ± *P_T_*. Within this range, the manager considers *N_t_* close enough to *T_N_*, and consequently, that the current policy results in a sustainable culling rate for the population. Hence, the manager will update the policy if and only if the population is monitored outside *T_N_* ± *P_T_*. The Trajectory (TRJ) strategy is the same as the ATI strategy, except that when *N_t_* is inside *T_N_* ± *P_T_*, the manager makes a prediction on next time step’s population size in the form of a linear extrapolation based on the current and preceding monitoring results. If this prediction falls inside *T_N_* ± *P_T_*, the manager leaves the policy unchanged; otherwise, they update it. In both ATI and TRJ strategies, after a time step without updating the policy, the manager receives an additional proportion *B_b_* of their initial budget to model the benefits associated with waiting. This bonus can be accumulated over several consecutive time steps of waiting but is lost as soon as the manager draws into their budget to raise the level of restrictions again (modelling details in Appendix 4).

#### Farmers action planning

At the beginning of each time step, each farmer receives a fixed, non-cumulative budget *B_F_*, which they allocate to culling a certain number of animals on the land that they own at the cost set by the manager’s policy. A minimum cost of 10 *b.u* models the baseline budget needed for a farmer to cull an animal. The number of animals culled is independently computed for each farmer using GMSE’s evolutionary algorithm, meaning that each farmer makes an independent decision for how to act according to their goal: maximizing their own yield.

### Simulations with GMSE

To simulate conservation conflict adaptive management with different timing strategies under uncertainty, we used the R package ‘GMSE’ (Duthie et al. 2018). See Appendix 4 for further details on modelling, parameter choices and simulations.

#### Initial parameters

We modelled the landscape as a grid of 40 equally sized rectangular pieces of land, each individually owned by a farmer. We model a population that is stable in absence of culling, but under an important threat of extinction for a high culling rate. We defined the population dynamics model parameters such that an equilibrium size was reached quickly and steadily, as a stable natural population would. The farmers were provided with an initial budget high enough to cull up to the expected number of animals on their land when population is at equilibrium (*B_F_* = 1000 b.u), so the population went extinct if the conflict is left unmanaged. At first, the manager’s initial budget was set equal to the farmers’ one (*B_M_* = *B_F_* = 1000 *b.u*) and manager’s target was set at half the equilibrium size (*T_N_* = 2000 animals). The initial population size was set to *N*_0_ = 1000 animals, which is sufficiently low for the population to be under immediate threat of extinction and justify the initial involvement of a manager. We intentionally chose these parameters for the Control strategy to produce adequate management while also leaving room for improvement in order to determine the extent to which alternative strategies can generate better results.

#### Population dynamics submodel

GMSE’s population dynamics model features a population of N animals, each of which has an age and a position on the landscape. In each time step, each animal moves from its current cell to a random cell within a defined range. Upon arrival, the animal consumes a proportion of 0.5 of the cell’s remaining yield. All animals move 12 times during a single time step in a random order. After all movement and feeding, animals asexually produce one offspring for every 5 resource units consumed, which are added to the population as new individuals. Next, animals that have consumed over 4.75 resource units and are younger than 5 time steps survive, the others are removed from the population. This consumption criteria leads to density-dependent intra-specific competition for resources, and modelling life events discretely generates inter-individual variability, as well as geographical and demographic stochasticity, therefore accounting for several sources of uncertainty around population dynamics.

#### Monitoring submodel

We assumed that the manager makes no errors during monitoring, thus *N_t_* represents the exact population size at each time step. This assumption avoided modelled stochastic monitoring errors that would have challenged a full understanding of management dynamics.

#### Decision-making submodel

In each time step, manager and farmer decision-making is independently modelled using evolutionary algorithms, allowing the emergence of a conflict when agents’ goals are opposed. This approach computes practical but not necessarily optimal decisions, recognizing that most people cannot think of every single possibility to choose the optimal one, but can choose the best option among those they could conceive (Hamblin 2012, S.I.1 in Duthie et al. 2018), generating uncertainty around stakeholders’ individual decision-making.

### Experimental plan

#### Systematic parameter exploration

To assess management quality of ATI and TRJ in terms of population dynamics and farmers yield, we varied the permissiveness (*P_T_*) and budget bonus (*B_b_*) parameters across a range of values for each strategy and compared the outcomes with those of CTL. *P_T_* ranged from 0% of the manager’s target (*T_N_*) (unconditional update at every time step, i.e., CTL) to 100% of *T_N_* (update only in the extreme situations where the population is extinct or close to natural equilibrium size) by 10% increments. *B_b_* ranged from 0% of the manager’s initial budget (*B_M_*) (no bonus following a time step of waiting) to 100% of *B_M_* by 10% increments. For each unique combination of *P_T_* and *B_b_*, we ran 100 independent simulation replicates of management over a period of 20 time steps under identical initial conditions.

#### Management outcomes

We defined the most desirable outcomes to be when management prevents the population from going extinct (1), while keeping it as close as possible to target (2) and ensuring the highest yield to farmers (3) with the lowest inequity among them (4). For a particular combination of parameters, extinction risk (1) was assessed as the frequency of extinction events over all replicates, denoted *f_ext_*. We measured how close to target the population was (2) with the difference between the population size (*N_t_*) and the manager’s target (*T_N_*) over *T_N_* at the end of a simulation averaged over all replicates, denoted *d_T_*, in % of *T_N_*. Farmers’ total yield (3) was calculated as the ratio of the sum of all cell’s yield at the end of a simulation over the maximum yield the landscape can provide in the absence of animal consumption (40000 yield units) averaged over all replicates and denoted *Y_end_* in % of the landscape’s maximum productivity. The among-farmer inequity (4) was measured as the difference between the lowest and highest farmer’s yields at the end of a simulation, averaged over all replicates, denoted *Y_ineq_*, in % of the highest yield. Finally, we computed the proportion of time steps without manager’s intervention over the time length of a simulation and averaged it over all replicates, denoted *t_w_*(1-*t_w_* is thus the proportion of policy updates). We computed 95% bootstrapped confidence interval around each average (Manly 2007). The between-stakeholder equity was assessed by systematically confronting the conservation and the agricultural outcomes to detect unbalanced repartition of costs and benefits.

#### Sensitivity to manager’s budget

We hypothesized that the effect of the budget bonus amount (*B_b_*) on management quality would be stronger in situations of higher budget constraint on the manager. To test for this, we selected the permissiveness of 50%, which outcomes with TRJ were not different from CTL but with a weak *B_b_* effect (see TRJ results section). We decreased the manager’s initial budget (*B_M_*) from 1000 to 500 *b.u* by 100 *b.u* increments. For each *B_M_*, we varied *B_b_* from 0 to 100% of *B_M_* by 10% increments in 100 replicates, and measured the same outcome proxies as the previous section to investigate the effect of *B_b_* amount on management quality according to *B_M_*. We also simulated management with CTL for each *B_M_* value in order to check how well the waiting strategies performed in comparison. This sums up to 60 different combinations of *B_M_* and *B_b_*, for an additional 6000 independent simulations.

## RESULTS

### Adaptive Timing of Intervention strategy

#### Conservation outcomes

When applying the Adaptive Timing of Intervention (ATI) strategy, increasing the permissiveness value caused the extinction risk to increase, and the final population size to decrease below target with no marked effect of the budget bonus (Figs. 1 and A1.1). No combination of permissiveness and bonus amount resulted in equivalent or lower extinction risk than CTL strategy (*f_ext_*= 0.15 with [0.08; 0.22] 95% confidence interval). No parameter combination of ATI strategy resulted in the population being closer nor equally close to target as CTL strategy (*d_T_* = −24.90% [−33.78; −16.26]) either, which is not surprising given that extinction was almost certain for most combinations (*f_ext_*> 0.9 for *P_T_*> 20%).

**Fig 1.**
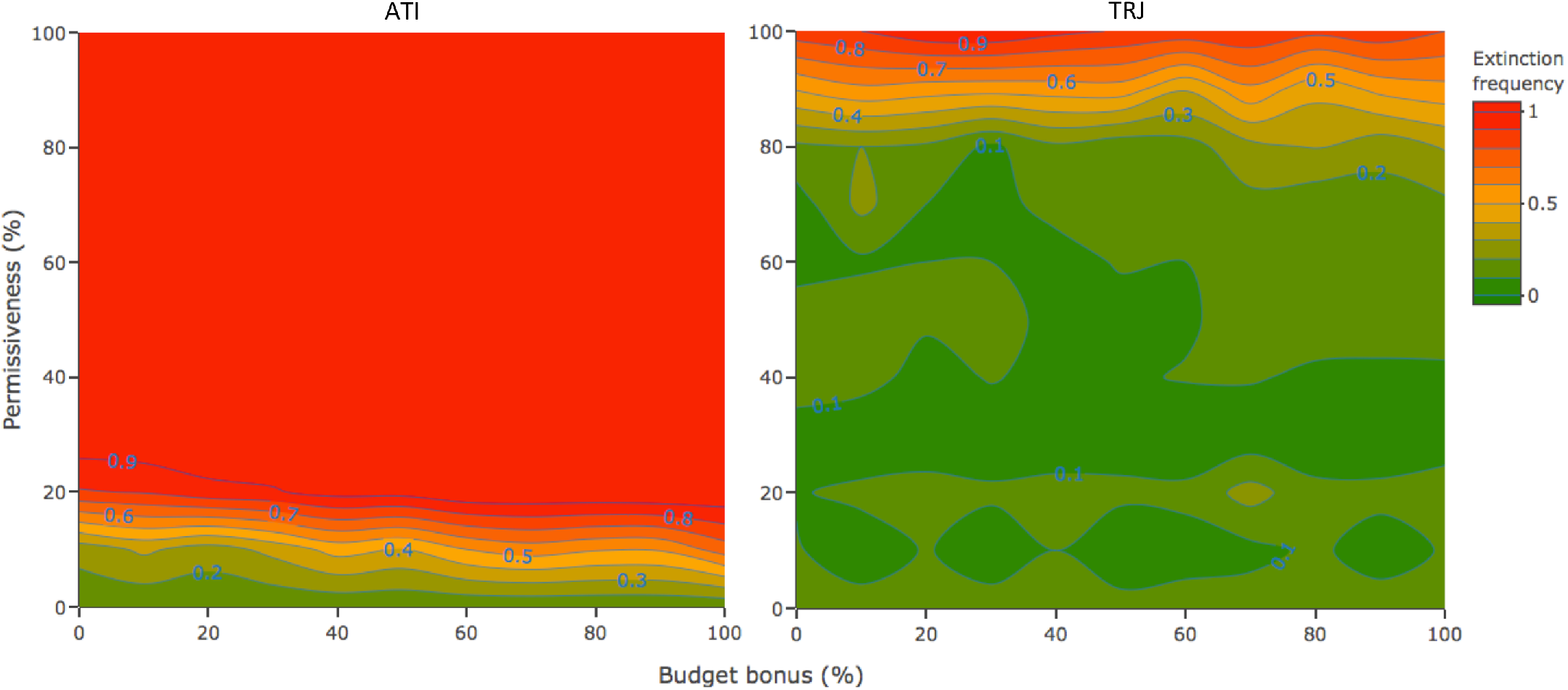
Extinction frequency (*f_ext_*) according to the permissiveness (*P_T_*) and budget bonus (*B_b_*) combinations in an individual-based model simulating the management of a population under conditions of conservation conflict. The *P_T_* = 0 and the corresponding *B_b_* values are the *f_ext_* obtained with the control strategy (CTL). The greener the lower the risk of extinction. With ATI (left panel), there was no combination of *P_T_* and *B_b_* parameters resulting in as low an fext as CTL (0.15 [0.08; 0.22] 95% CI), and population extinction was almost certain in most cases, with a weak positive effect of *B_b_* regardless of the permissive range size. With TRJ (right panel), most areas are as green as, to greener than CTL’s *f_ext_* value, meaning TRJ performed as least as good CTL regarding extinction risk. The effect of *B_b_* on *f_ext_* was weak to absent.

#### Agricultural outcomes

Increasing permissiveness caused the farmers’ final yield to increase, and among-farmer yield inequity to decrease with no effect of the budget bonus amount (Figs. A1.2 and A1.3). Farmers’ final yield was >90% of the maximum for all ATI parameter combinations, which was slightly more than CTL (*Y_end_*= 89.64% [88.04; 90.90]). The among-farmer inequity was slightly lower than CTL results (*Y_ineq_* = 5.68% [4.97; 6.34]). Indeed, as permissiveness increased, there were fewer animals feeding on farmers’ land so the impact on yield was lower and the farmers’ yield got closer to maximum. Also, the highest yields attained the maximum value while the lowest kept increasing, which reduced inequity.

#### Mechanisms underlying the outcomes

With ATI, most extinction events occurred when the population was monitored to exceed the permissive range, and in response, the manager lowered the level of culling restrictions to favor population decrease down to target. A problem arose when, in the following time step, the population was monitored inside the permissive range because it caused the manager to leave the policy unchanged. Farmers then continued to cull at a low cost, driving the population to extinction at the next time step (Fig. 2 ATI panel). Consequently, the larger the permissive range around target, the more likely this was to happen, thereby explaining why the extinction frequency and deviation from target increased with permissiveness values. This misinterpretation from the manager regularly occurred in the ATI parameter areas with very high extinction frequency (Fig. 1), in which the population deviation from target at the time step preceding extinction was within the manager’s permissive range (Fig. A1.4). Hence, the most effective strategy for avoiding population extinction here was to intervene unconditionally in every time step, at the expense of slightly decreasing farmers’ final yield.

**Fig 2.**
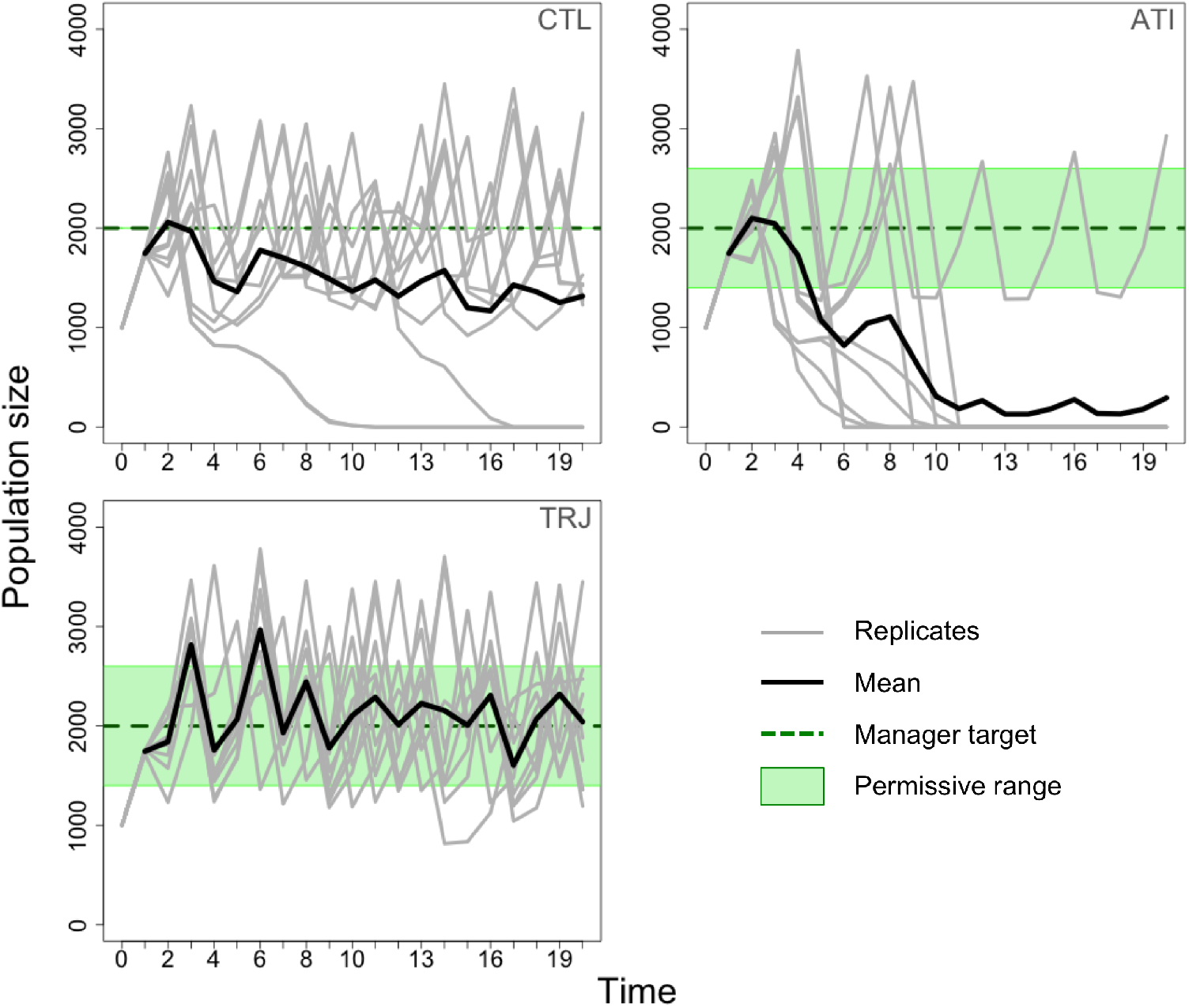
Average population size over time on ten simulation replicates with an individual-based model simulating the adaptive management of a population under conditions of conservation conflict. **Upper left**: when manager intervenes unconditionally (CTL). The extinctions happened when the population got too far below *T_N_* between two consecutive time steps for the manager to be able to rectify by increasing restrictions. **Upper right**: when applying ATI (*P_T_* = 30%, *B_b_* = 10%). Most extinctions happened when population size was over the permissive range, then was monitored into it the following time step. Thus, managers did not update the policy, allowing farmers to continue culling at a low cost, frequently driving the population to extinction at the following time step. Note that in the replicate that did not result in extinction, the population was never monitored into the permissive range during a decrease, causing the manager to update the costs and control the situation with better timing. Lower left: when applying TRJ (*T_N_* = 30%, *B_b_* = 0%). TRJ strategy avoided some extinction events.

### Trajectory strategy

#### Conservation outcomes

When applying TRJ, the extinction frequency and deviation from target were at least as close to 0 as CTL for permissiveness values up to 80%, without the manager intervening up to 40% of the time (Figs. 1, A2.1 and A2.2). The budget bonus value had either no effect or a weak effect on the outcomes. Several combinations resulted in an extinction frequency under 0.1, even 0 sometimes, while *f_ext_* = 0.15 [0.08; 0.22] with CTL. The effect of bonus amount was slightly stronger in the 40 and 50% permissiveness range (Fig. A2.2), where bonus values between 20 and 50% resulted in the population being closer to target than CTL (*d_T_* = −24.90% [−33,78; −16.26]). We chose the 50% parameter area for the experiment on sensitivity to manager’s initial budget to test whether this weak effect could amplify when applying stronger budget constraints on the manager.

#### Agricultural outcomes

With TRJ, the farmers’ final yield was as close to maximum, and the among-farmer yield inequity was similarly low as the CTL strategy regardless of the permissiveness and budget bonus value (Figs. A2.3 and A2.4).

#### Mechanisms underlying the outcomes

The rare extinction events with CTL seem to have occurred when population was over target and the manager decreased the level of restrictions by too much, or when farmers happened to cull more than expected, which caused the population to decrease beyond reparation (Fig. 2, CTL panel). TRJ strategy may have avoided this imprecision by offering managers the possibility not to intervene at these moments when the population is in the upper permissive range and keep the population closer to target (Fig. 2, TRJ panel). The absence of effect from the budget bonus amount was most likely caused by the manager initial budget alone often being enough to efficiently ensure both population maintenance and farmers’ yield given our initial parameter values, leaving no room for improvement due to a bonus. Thus, TRJ achieved similarly good management outcomes as CTL without managers having to intervene at every time step and regardless of the amount of benefit obtained from waiting periods.

### Sensitivity to manager’s initial budget

#### Conservation outcomes

The extinction frequency increased, and the final population size decreased below target with decreasing manager’s initial budgets (Fig. 3). But for *B_M_* = 800 b.u, the extinction frequency steadily decreased from 0.71 [0.61; 0.80] without budget bonus, to 0.07 [0.02; 0.12] for a bonus of 30% of B*M* (Fig. 3), which is significantly closer to zero than CTL for the same initial budget (*f_ext_* = 0.76 [0.67; 0.83]). At higher bonuses, the extinction frequency increased again between 0.3-0.6, which is lower than CTL, although still a high extinction risk. The same trend was observed in the distance to target, which rose from −78.4% of *T_N_* [84.9; −70.9] without budget bonus, to −11.4% [−21.4; −2.1] for the same bonus of 30% of *B_M_* (Fig. A3.1); CTL being −83.7% [−88.7; −78.0] (Fig. A3.1).

**Fig 3.**
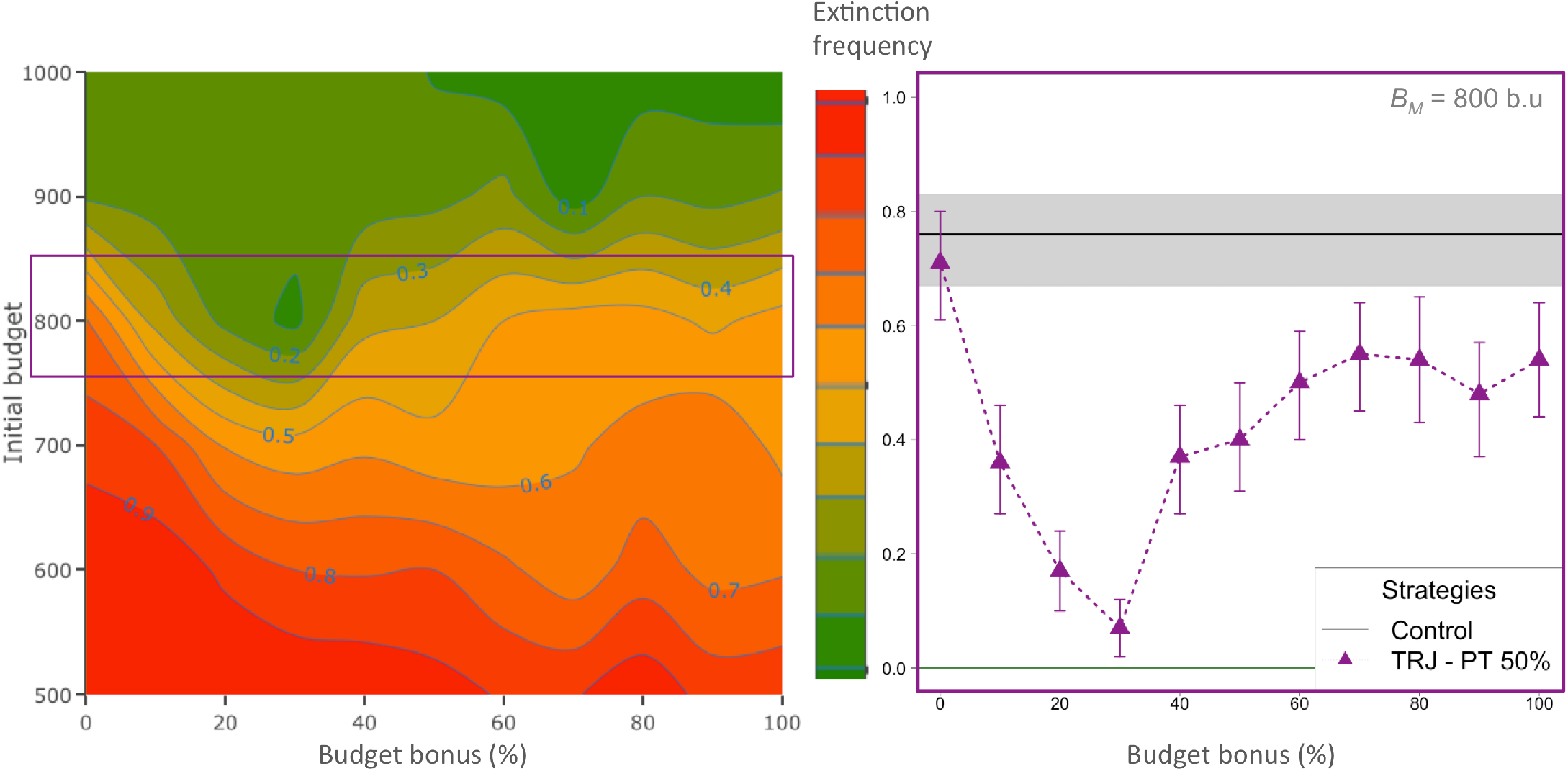
Extinction frequency when applying TRJ (*P_T_* = 50%) according to manager’s initial budget (*B_M_*) and budget bonus amount (*B_b_*) in an individual-based model simulating the adaptive management of a population under conditions of conservation conflict. The greener, the lower the extinction frequency. For *B_M_* = 800 *B_b_* (violet square, detail on the right panel), a pit forms along increasing *B_b_* values, meaning that low to intermediate values for *B_b_* markedly lowered the extinction risk. Error bars show 95% bootstrapped confidence intervals. The black line is the *f_ext_* with Control strategy for the same initial budget and the grey shaded area the 95% confidence interval around it.

#### Agricultural outcomes

The farmers’ final yield increased, and the among-farmer inequity decreased with decreasing manager’s initial budget (*B_M_*) because of the negative effect on extinction risk and population size (Figs. A3.3 and A3.4). In the *B_M_* = 800 *b.u* area, the farmers’ final yield was between 85% and 100% (for the highest extinction frequency) without varying markedly with the bonus amount. With the bonus of 30% that critically improved conservation outcomes, the final yield was 89.20% [87.47; 90.76] instead of 97.18% [96.14; 99] with the CTL strategy for the same manager’s budget (at the expense of a very high extinction risk). The inequity was 5.94% [5.23; 6.68] instead of 2.11% [1.65; 2.6] with CTL, which is still relatively low.

#### Mechanisms underlying the outcomes

For the manager’s initial budget value that maximized budget bonus’ negative effect on extinction risk and positive effect of on population size (*B_M_* = 800 *b.u*), if the manager intervened at every time step or used TRJ but without getting any benefit from the waiting periods, extinctions occurred when the population fell to too low a population size. At these moments, it was challenging for the manager to rectify the population trajectory with only their initial budget as the culling cost was always too low to efficiently reduce farmers culling rate (Fig. 4 CTL panel). If, in this situation, the manager accumulated budget bonus from previous waiting period(s), they had enough power to enforce higher restrictions to farmers as soon as the population did, or was predicted to, fall under the manager’s permissive range. Intermediate bonus amounts ensured that, when the latter happened, the population could increase closer to manager’s target (Fig. 4 TRJ panel). TRJ thus appeared to be more efficient than CTL in situations of stronger budget constraint on the manager. In such situations, the role of the budget bonus was critical in decreasing the extinction risk, while maintaining a high and equitable yield to farmers and allowing the manager to save 20 to 30% of their interventions (Fig. A3.5).

**Fig 4.**
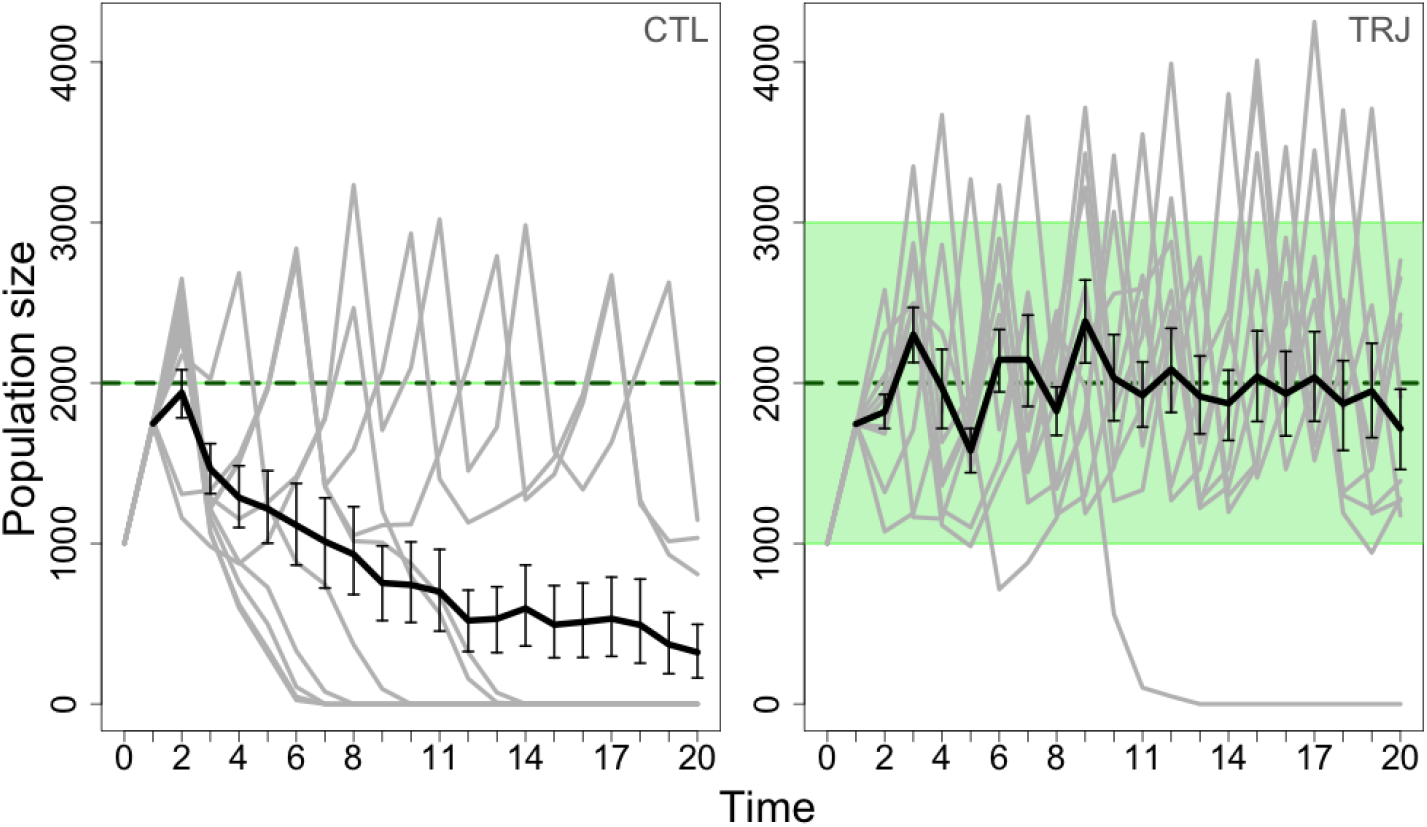
Population size over time averaged over 50 replicates (thick black line, error bars being the 95% confidence intervals) plotted on ten replicates (thin gray lines) with an individual-based model simulating the adaptive management of a population under conditions of conservation conflict and an initial budget of 800 *B_b_*. The green dotted line shows manager’s target *T_N_*, and the green area represents the permissive range *T_N_* ± *P_T_*. Left panel: when applying CTL. Extinctions happened when the population got too far below the manager’s target (green dotted line) between two consecutive time steps for the manager to be able to rectify with their initial budget only. Right panel: when applying TRJ (*P_T_* = 50%, *B_b_* = 30%). Thanks to the benefits accumulated over waiting periods, the manager was able to raise the culling cost high enough to maintain farmers’ culling rate at a sustainable value. The replicate that resulted in extinction was caused by a strong misprediction of time step 10’s population size, causing the manager to wait while intervention was needed.

## DISCUSSION

### Summary of the study

When adaptively managing a conservation conflict in a social-ecological system, our modelling of strategies dynamically alternating between intervention and waiting found that management outcomes were better when the decision to intervene was made based on a prediction of the system’s response than when based on the latest monitoring results alone. With prediction-based decisions, conservation and agricultural outcomes were at least as good as intervening unconditionally, while allowing the manager to save management resources and avoid unnecessary, potentially harmful interventions. When a low budget limited a manager’s ability to effectively manage the conservation conflict, the benefits accrued during waiting periods were applied when intervention was most critical and greatly improved conservation outcomes with only a weak impact on farmers’ yields and equity. Naturally, the main risk with waiting strategies is to decide to wait when intervention is needed, or to intervene when waiting is preferable. Basing intervention only on current monitoring should be avoided because when population density is monitored inside the permissive range during a sharp increase or decrease, managers can mistakenly conclude that the policy is adequate when, in fact, keeping the same policy running again can lead to extinction or critical yield loss. Basing intervention on population trajectory instead also includes a risk of inaccurately predicting the population density to be within the permissive range causing the managers to wait while the policy is inadequate to align conservation and agricultural objectives. Nevertheless, the consequences for yield loss or population decline were reversible when using an adequate permissive range.

### Importance of budget and monitoring in waiting strategy’s efficiency

The superiority of our Trajectory strategy over unconditional intervention depended on the manager’s budget. When the budget was high enough to manage the situation efficiently, the outcomes with the Trajectory strategy were at least as good as unconditional updates regardless of the budget bonus amount. This suggests that interventions when the population was monitored within the permissive range and predicted to stay in it (i.e., oscillating close to target) were less useful. Since the initial budget was sufficient for satisfactory management, the benefits reaped during waiting periods with the Trajectory strategy could not further improve the management outcomes. This is relevant because human, financial and time resources are limited in conservation and there is a constant competition for their allocation to cases (Hughey et al. 2003, Jachowski and Kesler 2008, McDonald-Madden et al. 2008, Ruiz-Miranda et al. 2020). It is also increasingly recognized that different species can impact human livelihood in different ways and at different times within the same geographical area, which should be considered in management (Pozo et al. 2020, 2021). Intervention in one conflict could thus be a priority for a time, and then deprioritized when another requires intervention more urgently. Therefore, resources unused during periods of waiting in a well-funded case could instead be allocated to other, potentially less well-funded and/or more pressing cases and improve overall conservation benefits (Wu et al. 2020). Our Trajectory strategy can thus help a dynamic allocation of management resources to cases that need them the most at a given instance.

When a limited budget made management more challenging, the resources saved when not intervening using the Trajectory strategy could generate enough benefits to compensate for the lack of resources. We emphasize that the prediction based on population trajectory is a means for managers to reduce the risk of misjudging the timing of intervention; what improved management here was better access to the benefits accumulated over waiting periods. This result supports previous modelling results in Iacona et al. (2017), where national park managers did not have enough budget to put every endangered bird species under protection at once but could maximize success by waiting and saving their funds to gradually enhance their monetary power. Importantly, this is only possible if unused management resources are not revoked or reallocated when less needed. A review of exit-strategies in conservation by Ruiz-Miranda et al. (2020) found that withdrawing funds when objectives are attained is very uncommon in adaptive management (but should be more considered and carefully planned). The present study suggests that the budget saved during waiting periods should be reallocated if the management resources are not limiting but invested in improving future interventions if they are.

To isolate the effect of various timing strategies on management quality, we assumed that the manager had perfect knowledge of population size. But real-world monitoring involves uncertainty that plays an important role in the success of conservation (Bunnefeld et al. 2011, Nuno et al. 2013). Monitoring uncertainty will cause errors in estimating population density, and therefore errors in deciding if the situation requires intervention. This will decrease the efficiency of both unconditional intervention and Trajectory strategies, but the latter might be more impacted because errors will influence both monitoring and trajectory prediction, therefore mitigating the advantage over unconditional intervention. Indeed, the efficacy of Trajectory strategy might rely on more regular and accurate monitoring, which might not always be possible or affordable. Testing the effect of observation accuracy or cost on management quality is beyond the scope of this study, but it is an important aspect to consider when applying timing strategies (Milner-Gulland 2011, McDonald-Madden et al. 2011, Wu et al. 2020).

Since our focus is on management strategy and not on control measures, we limited farmers’ options to culling for the sake of simplicity and ease of model interpretation. We did not model indirect measures such as fencing, widespread in the management of conservation conflicts over land-use (Nyhus 2016, Pooley et al. 2017), as these measures are rather permanent constructions that are not always fitted to the regular changes and updates of our adaptive management process. Nevertheless, future modelling might usefully consider a range of alternative options for population management.

### Modelling novelties for adaptive management

The ongoing 6th mass extinction under a rapidly changing climate (Ceballos et al. 2017) and the consequences of land-use conflicts between agriculture and wildlife protection on food security often put conservation managers under urgency (Du Toit 2010). Our results suggest that the urgency to act should not mean systematic, unconditional intervention and stress the importance of acquiring information to choose wisely how and when to intervene. As with software such as ISIS-fish (Mahévas and Pelletier 2004) or FLR (Kell et al. 2007) in fisheries management, the method developed here can inform managers’ policy-making. Parametrizing GMSE with empirical data from a conflict between farming and common cranes in Sweden has previously permitted the evaluation of subsidy levels that best balanced culling and scaring for the maintain of both population and farmers’ income (Nilsson et al. 2020). Likewise, targeted parametrization of our model can give managers information to decide how permissive they should be and how much gain they should expect from waiting periods for our strategy to be useful regarding conservation and land-users’ objectives, management resources allocation efficiency.

The individual-based nature of our model and the modularity of the GMSE framework accounts for several sources of uncertainty around population dynamics and stakeholders’ individual decision making. Our mechanistic model simulates population dynamics with intrinsic demographical uncertainty (inter-individual variability in the realization of life events) and geographical uncertainty (animals’ movement is stochastic; Uchmański and Grimm 1996; Stillman et al. 2015). Future work could also include explicit modelling of environmental uncertainty, potentially in the form of stochastic extreme events impacting both population dynamics and farmers’ yields. Currently, our results are robust even if population dynamics are uncertain and if spatial distribution can induce inequity by having the animals sometimes being more numerous on one farmer’s land than another. Yet, Rakotonarivo et al. (2020, 2021) showed that the perceived equity in the balance of costs and benefits of conservation actions between and among stakeholders’ groups plays an important role in land-users’ propensity to choose pro-conservation strategies. However, the aspect of equity in conservation conflicts has scarcely been incorporated in modeling results. For example, Wam et al. (2016) used a measure of monetary equity between different stakeholder groups in their management model balancing logging, livestock grazing and game hunting activities in a boreal forest. Our method also controls between-stakeholder equity by systematically confronting the population dynamics and the farmers’ yield. In addition, we used a new indicator parameter for among-stakeholder equity by measuring the success of our strategies against the difference between the lowest and highest farmers’ yields. Among-stakeholder equity, to our knowledge, has not been modelled before in conservation conflicts, and modelling stakeholders individually like the present study offers a direct measure of equity among members of the same group, thus allowing its monitoring as an important outcome of management.

The lack of dynamic stakeholder behavior modelling has been identified as a major cause of failure in conservation (Schlüter et al. 2011). Previous studies have addressed this by modelling decision-making using game theory (Colyvan et al. 2011, Glynasty et al. 2018). Nevertheless, a game-theoretic framework can have limitations when applied to management decision-making, including fixed behavior rules, finite sets of actions (e.g., cooperate or defect) and the assumption that players are perfectly rational and aware of the best options for them (Myerson 1997). In this model, we use evolutionary algorithms, a form of artificial intelligence, for managers and farmers to make decisions, which we show here offers a heuristic to find practical solutions when the panel of options is too large for game theoretic problems (Hamblin 2012). We combined the evolutionary algorithms with an individual-based approach and model decision-making independently for each stakeholder with the possibility for sub-optimal choices along a continuum of possible actions (see also Kamra et al. 2018, Cusack et al. 2020, Nilsson et al. 2021). Simulating these different sources of uncertainty in our experiments allowed to conclude that the strategy we proposed is relevant even if managers do not always make the most efficient policies and if farmers do not always behave as they were expected to.

## Conclusion

We use an uncertainty-robust modelling tool to compare the management quality of waiting strategies against unconditional intervention regarding conservation and agricultural objectives and discuss which strategy to prefer according to cases of conservation conflicts. We propose a strategy for managers to dynamically alternate between intervening and waiting informed by population monitoring. When the decision to intervene or wait is based on a prediction of population trajectory, our strategy can result in a better, more equitable management of conservation conflicts, especially in situations of limiting budget. By saving time, energy and/or money when intervention is not necessary, it can also ensure a more efficient use of management resources.

## Supporting information

ATI experiment's extended results

TRJ experiment's extended results

TRJ sensitivity to manager's budget experiment's extended results

Modelling details

French version of the abstract

## Acknowledgments

A.B. was funded by IAPETUS NPIF allocation, grant code NE/R012253/1 and supported by NERC, A.B.D. was funded by a Leverhulme Trust Early Career Fellowship, J.M. and N.B. were funded by the European Research Council under the European Union’s H2020/ERC grant agreement no 679651 (ConFooBio).

## Notes

ttps://github.com/AdrianBach/EcologyAndSocietyArticle-data-code.git

### Competing Interest Statement

The authors have declared no competing interest.

### Summary of Updates

This version has been revised according to two anonymous reviewers chosen by the journal Ecology and Society's editorial board. We made the statements about the management benefits associated with waiting more precise and referenced. We condensed the manuscript by removing paragraphs that were redundant between sections, and lowered the level of modelling details in the methods (all the details in Appendix 4). In discussion, we reminded that our example was focused on culling and that our results might be different for other control method. We also expanded on the implications of our work for adaptive management, particularly in the context of environmental urgency.

