## Supplementary material for "Intervene or wait? Modelling the timing of intervention in conservation conflicts adaptive management under uncertainty": ATI experiment's extended results

**Appendix 1.**

Additional figures of the adaptive timing of intervention strategy experiment results.

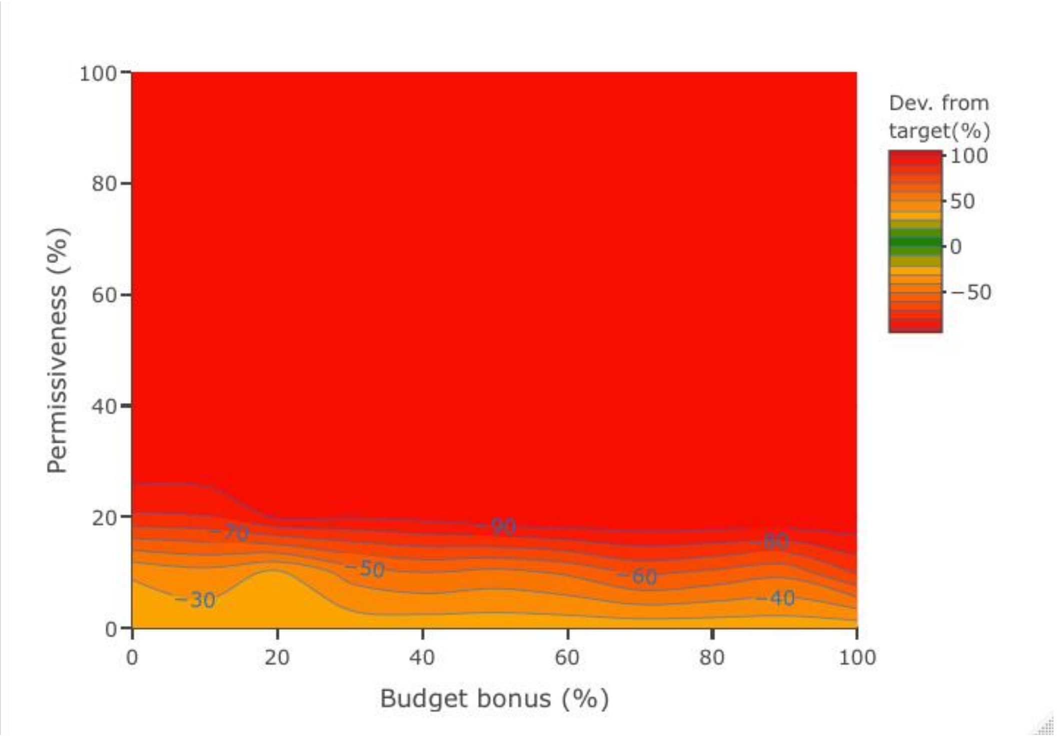


Figure A1.1. Population's average deviation from target ($d_T$) at the final time step of simulation according to permissiveness ($P_T$) and budget bonus ($B_b$) values when applying the Adaptive Timing of Intervention strategy. Results from simulations with an individual-based model simulating the adaptive management of a population under conditions of conservation conflict. The greener, the closer the population to manager's target ($T_N$). Given the numerous extinctions (see Fig.1), the population very often ended at a size of 0, meaning a –100% deviation from target, hence the large red area. With control strategy, the population was under target by –30 to –20%. Expectedly, this reflects the same tendency as control strategy.

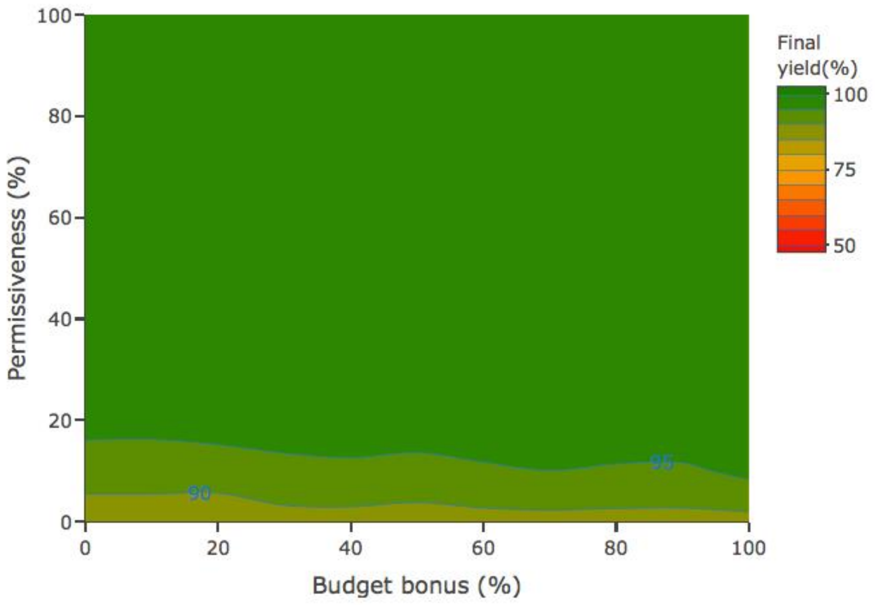


Figure A1.2. Average farmers' yield ($Y_{end}$) at the final time step of simulation according to permissiveness ($P_T$) and budget bonus ($B_b$) values when applying the Adaptive Timing of Intervention strategy. Results from simulations with an individual-based model simulating the adaptive management of a population under conditions of conservation conflict. The greener, the closer the farmers' yield to landscape maximal productivity. Given the numerous extinctions (see Fig.1), farmers very often reach their maximal yield, hence the large green area. With control strategy, farmers got between 85 and 90% of their maximal yield in average because the population was more efficiently managed and thus larger.

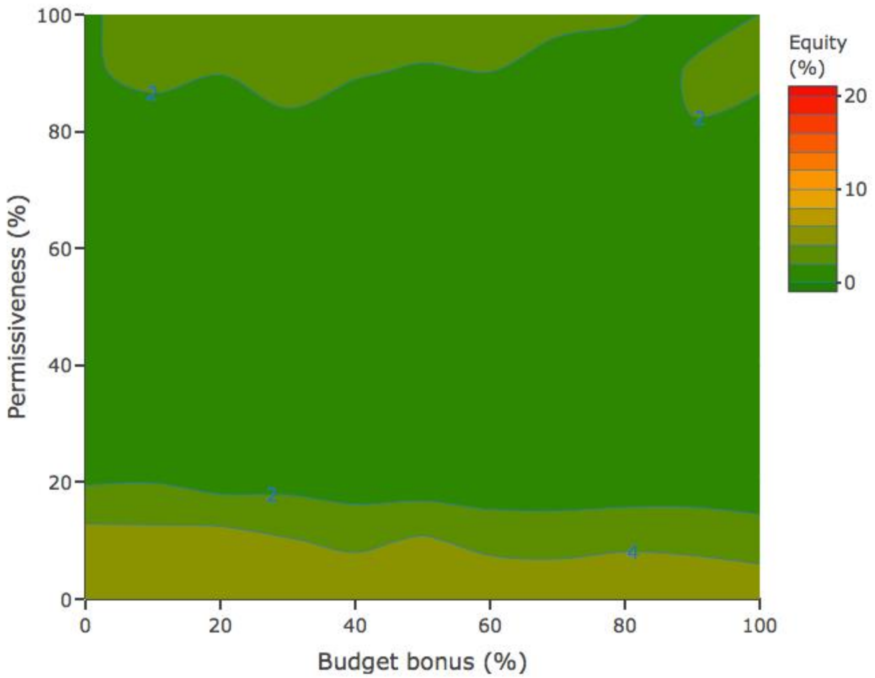


Figure A1.3. Average farmers' yield inequity ($Y_{ineq}$) at the final time step of simulation according to permissiveness ($P_T$) and budget bonus ($B_b$) values when applying the Adaptive Timing of Intervention strategy. Results from simulations with an individual-based model simulating the adaptive management of a population under conditions of conservation conflict. The greener, the smaller the difference between the highest and lowest farmer's yields. Given the numerous extinctions (see Fig.1), farmers very often reach their maximal yield while the lower yields were higher than with control strategy, hence the very low inequity.

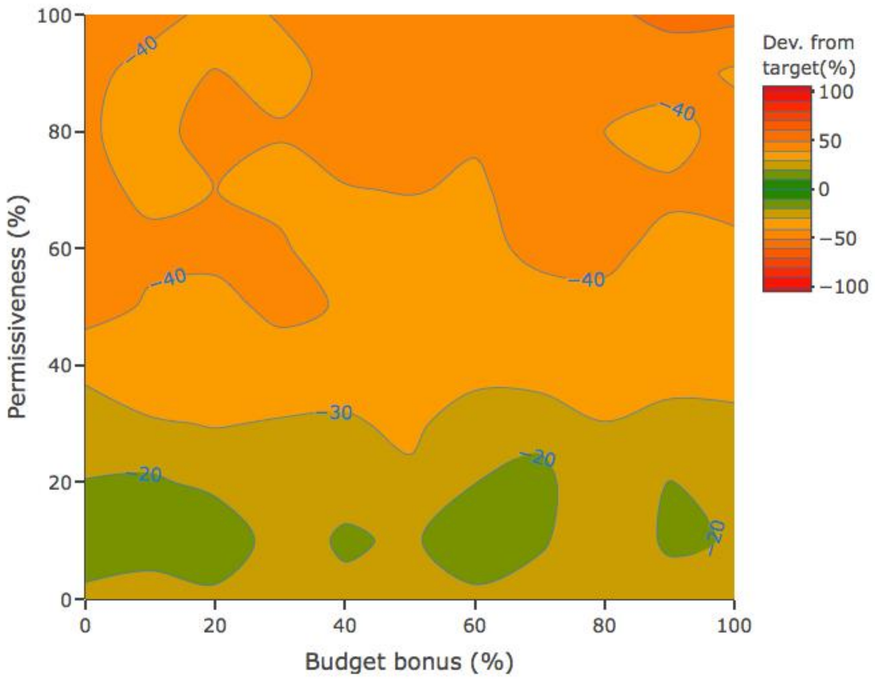


Figure A1.4. Population's average deviation from target ($d_T$) at the time step before the end of simulation ($t_{max}$ or extinction) according to permissiveness ($P_T$) and budget bonus ($B_b$) values. Results from simulations with an individual-based model simulating the adaptive management of a population under conditions of conservation conflict. The greener, the closer the population to manager's target ($T_N$). Note that in most areas of high extinction risk (red areas in Fig.1), the population size was monitored into the corresponding permissive range in the time step preceding extinction, causing the manager to wait when intervention was urgent.

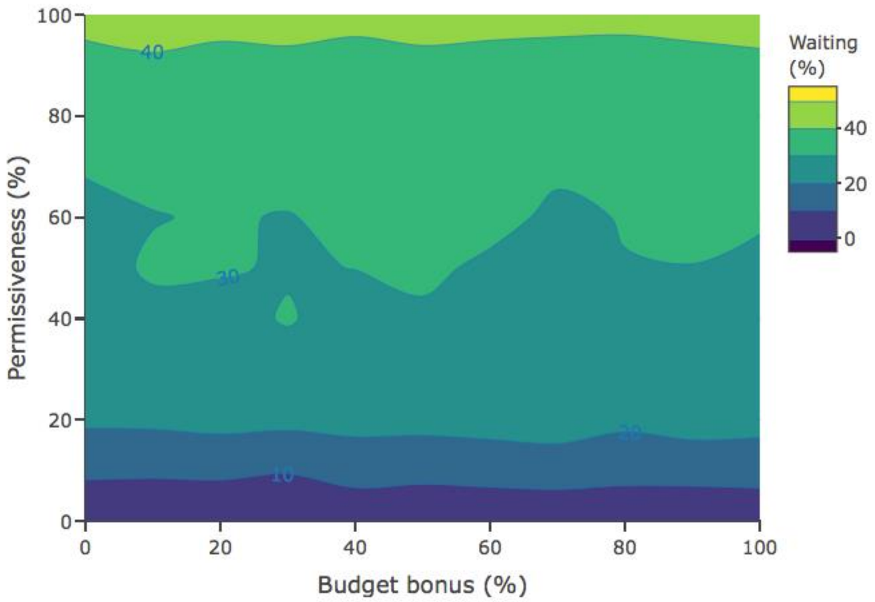


Figure A1.5. Average proportion of time steps without manager's intervention ($t_w$) during a simulation according to permissiveness ($P_T$) and budget bonus ($B_b$) values when applying the adaptive timing of intervention strategy. Results from simulations with an individual-based model simulating the adaptive management of a population under conditions of conservation conflict. The lighter, the larger the number of time steps without intervention.
