## Supplementary material for "Intervene or wait? Modelling the timing of intervention in conservation conflicts adaptive management under uncertainty": TRJ experiment's extended results

**Appendix 2.**

Additional figures of the trajectory strategy experiment results.

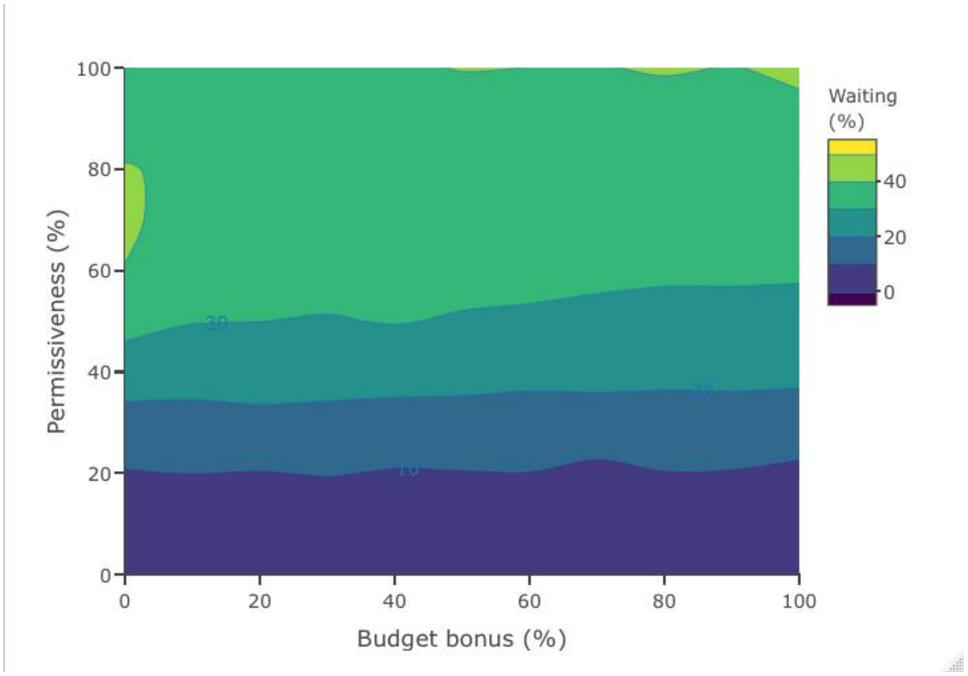


Figure A2.1. Average proportion of time steps without manager's intervention ($t_w$) during a simulation according to permissiveness ($P_T$) and budget bonus ($B_b$) values when applying the Trajectory strategy. Results from simulations with an individual-based model simulating the adaptive management of a population under conditions of conservation conflict. The lighter, the larger the number of time steps without intervention. In the 30% $P_T$ parameter area, the manager could save between 10 and 20% of their interventions.

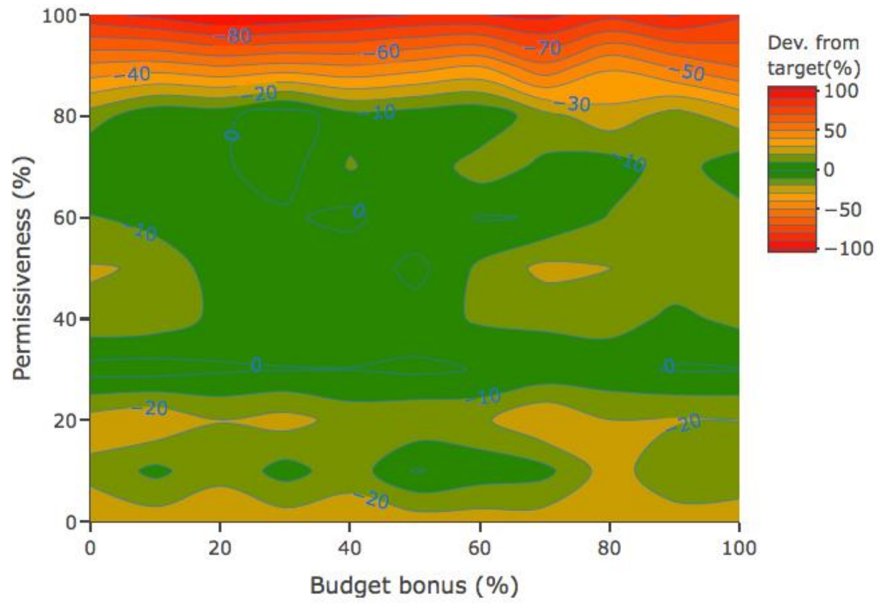


Figure A2.2. Population's average deviation from target ($d_T$) at the final time step of simulation according to permissiveness ($P_T$) and budget bonus ($B_b$) values when applying the Trajectory strategy. Results from simulations with an individual-based model simulating the adaptive management of a population under conditions of conservation conflict. The greener, the closer the population to manager's target ($T_N$). Most areas are greener than the control strategy ($P_T$ = 0 band) meaning that the trajectory strategy maintained the population closer to target. Note that in the $P_T$ = 30 parameter area, $d_T$ is the closest to 0 for every $B_b$ values.

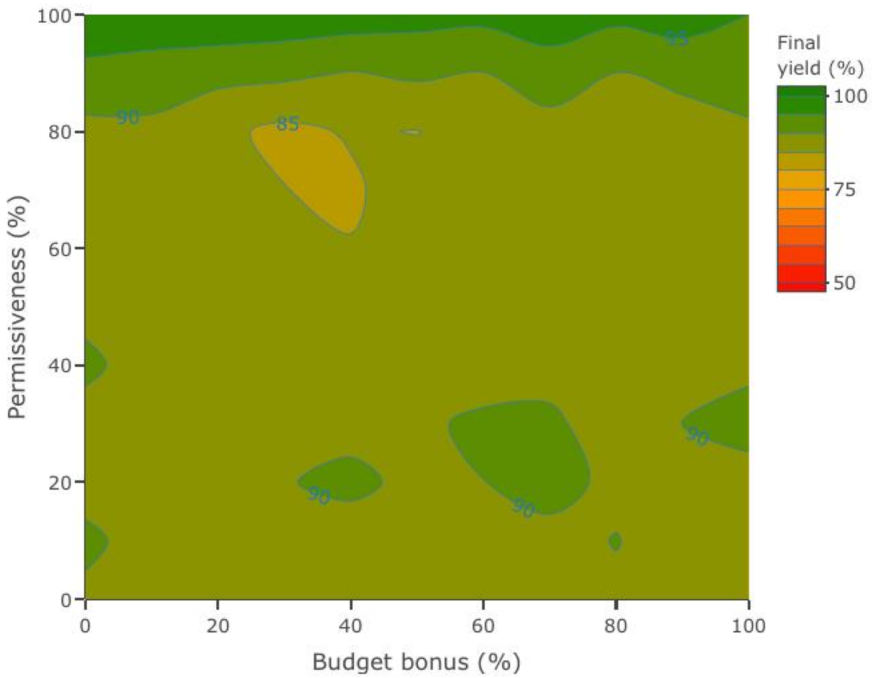


Figure A2.3. Average farmers' yield ($Y_{end}$) at the final time step of simulation according to permissiveness ($P_T$) and budget bonus ($B_b$) values when applying the Trajectory strategy. Results from simulations with an individual-based model simulating the adaptive management of a population under conditions of conservation conflict. The greener, the closer the farmers' yield to landscape maximal productivity. Most areas are as green as control strategy, with a final farmers' yield over 85% of their maximum.

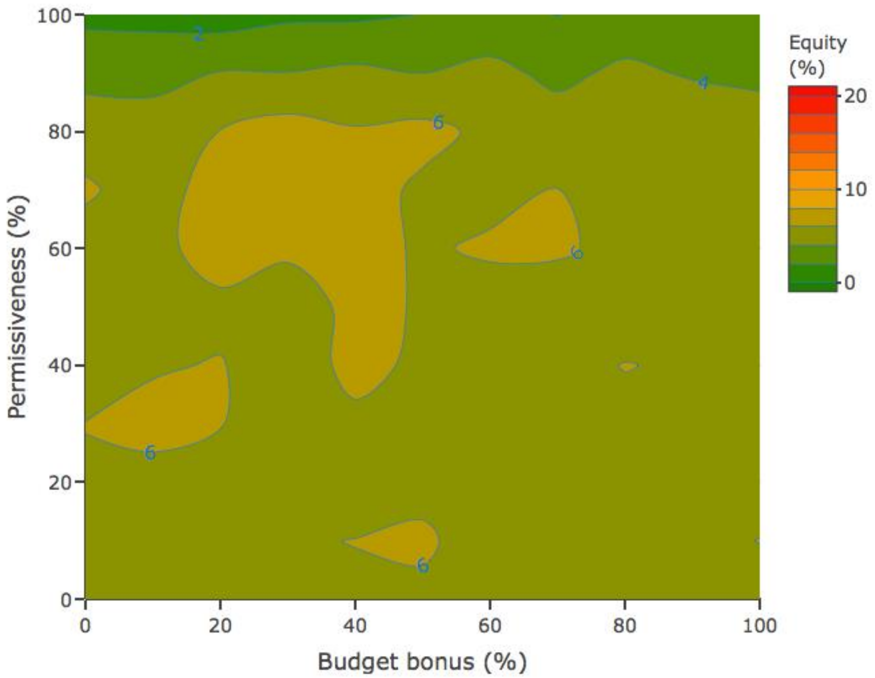


Figure A2.4. Average farmers' yield inequity ($Y_{ineq}$) at the final time step of simulation according to permissiveness ($P_T$) and budget bonus ($B_b$) values when applying the trajectory strategy. Results from simulations with an individual-based model simulating the adaptive management of a population under conditions of conservation conflict. The greener, the smaller the difference between the highest and lowest farmer's yields. Most areas are as equitable, or slightly less equitable than control strategy.
