## Supplementary material for "Intervene or wait? Modelling the timing of intervention in conservation conflicts adaptive management under uncertainty": TRJ sensitivity to manager's budget experiment's extended results

**Appendix 3.**

Additional figures of the sensitivity to manager's initial budget experiment results.

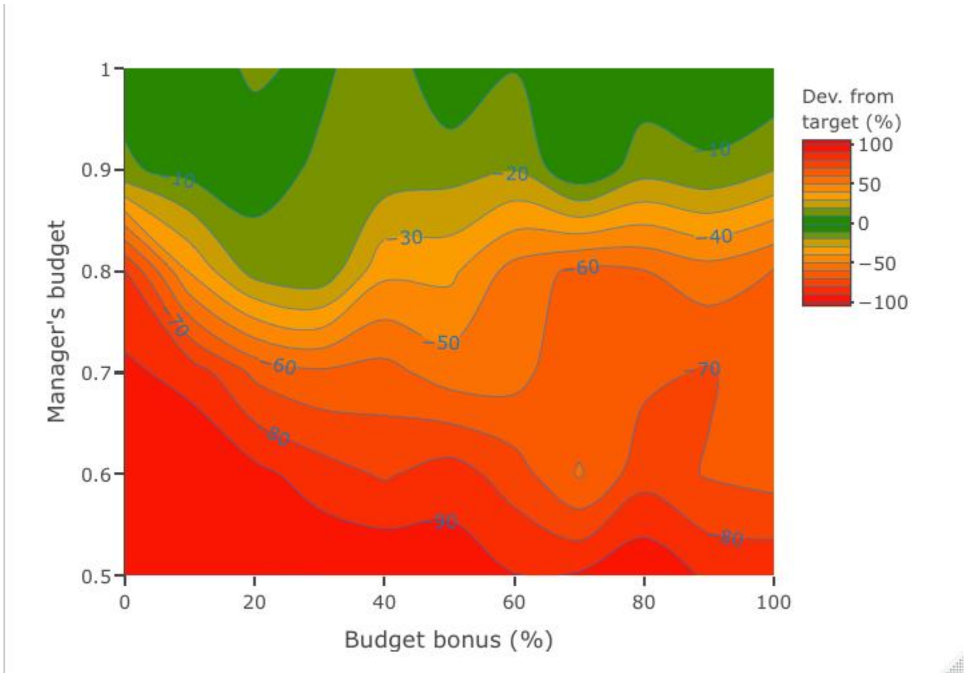


Figure A3.1. Population's average deviation from target ($d_T$) at the final time step of simulation according to manager's initial budget ($B_M$) and budget bonus ($B_b$) values when applying the Trajectory strategy. Results from simulations with an individual-based model simulating the adaptive management of a population under conditions of conservation conflict. The greener, the closer the population to manager's target ($T_N$).

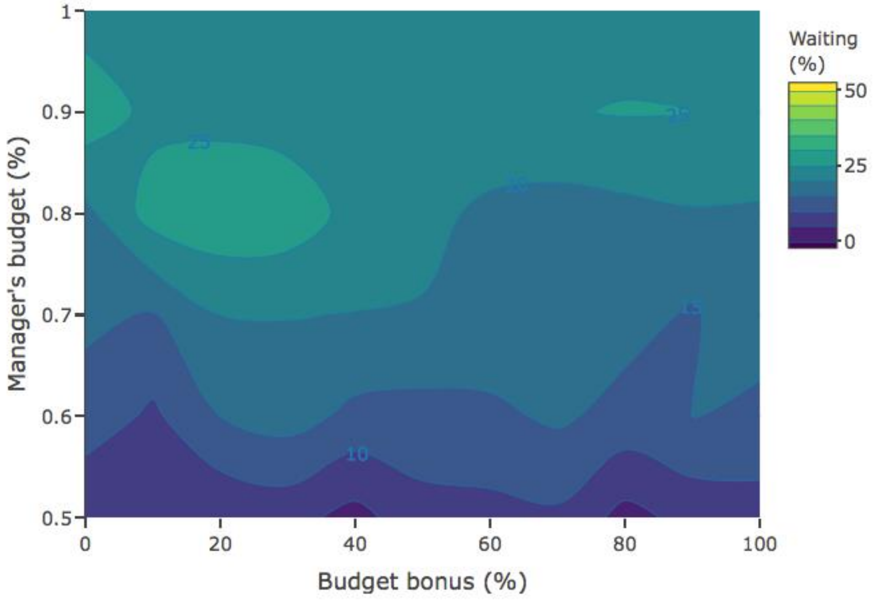


Figure A3.2. Average proportion of time steps without manager's intervention ($t_w$) during a simulation according to manager's initial budget ($B_M$) and budget bonus ($B_b$) values when applying the Trajectory strategy. Results from simulations with an individual-based model simulating the adaptive management of a population under conditions of conservation conflict. The lighter, the larger the number of time steps without intervention. The $B_M$ = 800 b.u. and 20-30% $B_b$ parameter area was also the one where the manager needed to intervene less, another sign that the population is often close enough to target not to need an intervention.

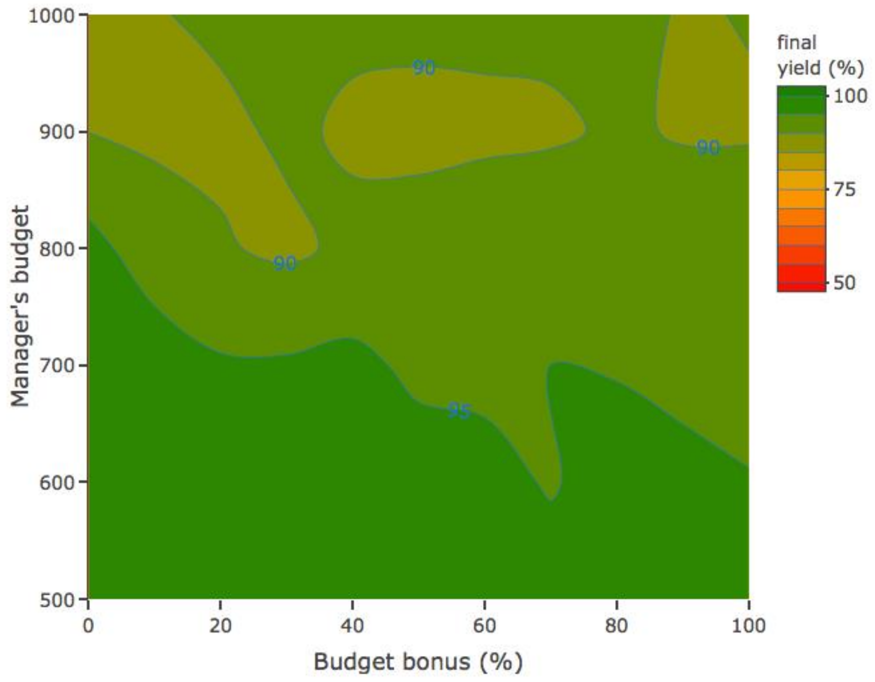


Figure A3.3. Average farmers' yield ($Y_{end}$) at the final time step of simulation according to manager's initial budget ($B_M$) and budget bonus ($B_b$) values when applying the Trajectory strategy. Results from simulations with an individual-based model simulating the adaptive management of a population under conditions of conservation conflict. The greener, the closer the farmers' yield to landscape maximal productivity. In the areas where the extinction frequency is acceptable, the farmers' final yield over 85% of their maximum, which is comparable to the previous experiments.

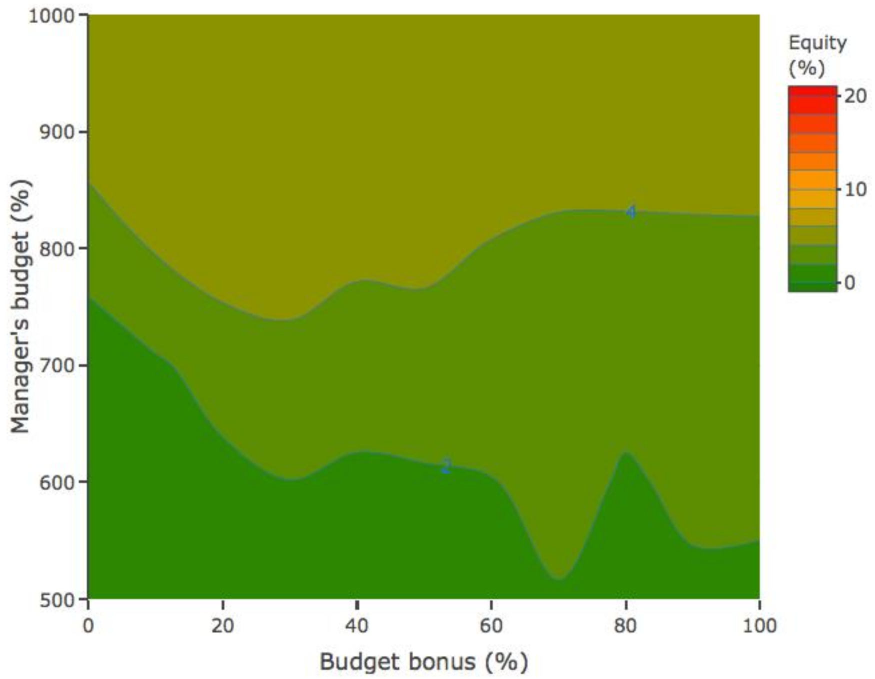


Figure A3.4. Average farmers' yield inequity ($Y_{ineq}$) at the final time step of simulation according to manager's initial budget ($B_M$) and budget bonus ($B_b$) values when applying the trajectory strategy. Results from simulations with an individual-based model simulating the adaptive management of a population under conditions of conservation conflict. The greener, the smaller the difference between the highest and lowest farmer's yields. In the areas where the extinction frequency is acceptable, the inequity is between 4 and 6% which is comparable to the previous experiments.
