## Supplementary material for "Intervene or wait? Modelling the timing of intervention in conservation conflicts adaptive management under uncertainty": Modelling details

**Appendix 4.**
Modelling details.

### Model overview

##### *Model case.*

To simulate conservation conflict management over time, we develop an individual-based model with a population of discrete animals, discrete farmers, and a biodiversity manager, all interacting on an agricultural landscape. The landscape is divided into discrete cells, each of which produces an agricultural yield and can hold any number of animals. Each farmer owns a contiguous block of cells that forms their 'land', and the sum of its cells' productivity determines the farmer's yield. Each animal's reproduction and survival depend on the amount of agricultural resources it consumes from landscape cells, which consequently reduces the farmers' yield. Farmers can cull animals that are on their own land to reduce yield loss. We chose population parameter values to ensure that unrestricted culling consistently drove the animal population to extinction (see the 'initial parameters' section below). The manager attempts to avoid extinction by maintaining the population around a predefined target size ($T_N$). This target was chosen to be high enough to prevent extinction, but low enough to ensure a satisfactory yield to farmers. The manager's method is to implement a policy incentivizing or disincentivizing culling as appropriate to increase or decrease population size to be closer to $T_N$. Hence, following an adaptive management process, the manager updates this policy according to the monitoring of the population size ($N_t$) at each time step $t$. Farmers' and manager's actions are constrained by finite budgets (respectively $B_F$ and $B_M$), which we interpret to reflect the total time, energy or money that a farmer can allocate to realize culling actions, or the manager to implement a change of policy and enforce culling restrictions at each time step. Furthermore, a conservation conflict will arise when the policy enforced by the manager prevents the farmers from culling as many animals as they want to minimize yield loss. Our case's conflict dynamics are therefore affected by both the ecology of the population and the flexible, goal-oriented decision-making of the manager and farmers.

##### *Manager policymaking.*

To maintain the population as close as possible $T_N$, the manager receives a fixed, non-cumulative budget $B_M$ at the beginning of each time step (i.e., it is completely lost if unused at the end of the time step). They can allocate it into setting a cost that farmers must pay to cull an animal on their land. A minimum cost of 10 budget units (b.u.) models the baseline budget needed for a farmer to cull an animal. The manager can draw into $B_M$ to raise this cost to discourage farmers from culling and favor population growth and can decrease it to facilitate culling and favor a population decrease. To model the budget needed to enforce a policy restricting culling, a raise of 1 in the culling cost requires an investment of 10 b.u. from the manager. Conversely, as the manager does not need to incentivize farmers to remove animals when the policy allows high culling rates, they do not need to spend budget to decrease the cost. The amount by which the manager changes the culling cost is computed according to their goal (see the 'decision-making sub-model' section below), i.e., keeping the population as close as possible to target. Manager's goal was modelled as minimizing the distance between the monitored population size $N_t$ and $T_N$.

##### *Timing strategies.*

We included three timing strategies that determine whether a manager intervenes and updates the policy or waits and leaves it as is. The Control strategy (CTL) was the null model in this study. It corresponds to unconditional intervention at every opportunity and was modelled as the manager simply updating the policy at every time step. With the Adaptive Timing of Intervention strategy (ATI), the manager dynamically alternates between intervening and waiting based on the distance between $N_t$ and $T_N$. ATI defines a permissive range $P_T$ around $T_N$ in the form of $T_N \pm P_T$. Within this range, the manager considers $N_t$ close enough to $T_N$, and consequently, that the current policy results in a sustainable culling rate for the population. Hence, at a given time step, the manager will update the policy if and only if the population is monitored outside this $T_N \pm P_T$ range. The Trajectory (TRJ) strategy is the same as the ATI strategy, except that when $N_t$ is into $T_N \pm P_T$, the manager makes a prediction on next time step's population size based on the current and preceding monitoring results. If this prediction falls into the $T_N \pm P_T$ range, the manager assumes that the policy is effective and leaves it unchanged; otherwise, they update it. In both ATI and TRJ strategies, after a time step without updating the policy, the manager receives an additional proportion $B_b$ of $B_M$ to model the benefits associated with waiting (e.g., the money, time or energy saved by not engaging in the process of updating the policy and enforce the change on farmers, or the interests gained from putting up the money saved). This bonus can be accumulated over several consecutive time steps of waiting but is lost as soon as the manager draws into their budget to raise the level of restrictions again.

##### *Farmers action planning.*

At the beginning of each time step, each farmer receives a fixed, non-cumulative budget $B_F$, which they allocate into culling a certain number of animals on the land that they own at the cost set by the manager's policy. The number of animal culled is independently computed for each farmer using GMSE's evolutionary algorithm (see the 'decision-making sub-model' section below), meaning that each farmer makes an independent decision for how to act according to their goal: maximizing their own yield. We used this model case to investigate how different timing strategies for a biodiversity manager's intervention can affect the outcomes of an adaptively managed conservation conflict.

#### Simulations with GMSE

To simulate a conservation conflict management with different strategies under uncertainty, we used the R package 'GMSE' (Duthie et al. 2018). GMSE is a flexible modelling tool to simulate key aspects of natural resource management over time and address adaptive management questions *in silico* (Cusack et al. 2020, Nilsson et al. 2021). GMSE offers a range of parameters to simulate resource variations and management policy options with individual-based models of population dynamics, monitoring, manager decision-making and farmer decision-making.

##### *Initial parameters.*

We modelled a spatially explicit landscape with a grid of 200 by 200 cells, divided into 40 equally sized rectangular pieces of land, each individually owned by one of 40 farmers. For the animals, we wanted to model a population that is stable in absence of culling, but under an important threat of extinction under a high culling rate. We defined the population dynamics model parameters such that, under constraint of density-dependent intra-specific resource competition only, an equilibrium was reached quickly and steadily, as a stable natural population would. The size at equilibrium ($K$) was sought such that the expected number of animals per farmer's land was about a hundred on average (i.e., around 4000 individuals on the landscape). The farmers were provided with an initial budget high enough to cull up to the expected number of animals on their land at the baseline cost (i.e., 1000 b.u), and at first, the manager's initial budget was set equal to the farmers' one. We set $T_N$ at half the equilibrium size, which was low enough to maintain farmers' yield over 90% of their maximum yield, but high enough to ensure a relatively low extinction risk of around 15% with the Control strategy (c.f. Management outcomes and Results sections in main document). We intentionally chose these parameters for the Control strategy to produce adequate management while also leaving room for improvement in order to determine the extent to which alternative strategies can generate better results. We set the initial population size $N_0$ = 1000, which is sufficiently far below $K$ for the population to be under extinction threat and justify the initial involvement of a manager.

##### *Population dynamics sub-model.*

GMSE's population dynamics model features a population of $N$ animals, each of which has an age as well as an $x$ and $y$ landscape position, all initialized at random (integers sampled with equal probabilities along the range of possible values). In each time step, each animal moves from its current cell to a random cell within a defined range of cells in any direction (including the original cell). After arriving at a cell, the animal feeds and consumes a proportion of 0.5 of the cell's remaining yield. All animals move 12 times during a single time step, but individual movement across all animals occurs in a random order to avoid having a subset of animals complete all their moving and feeding before the others have started. After all movement and feeding has occurred, the animals asexually produce one offspring for every 5 resource units consumed (e.g., if an animal has consumed 12 resource units it produces 2 offspring). The offspring are added to the population as new individuals of age 0 on the cell on which they were produced. Next, animals that have consumed over 4.75 resource units and have an age under or equal to 5 time steps survive to the next one. Animals that do not survive are removed from the population. This consumption criteria lead to density-dependent intra-specific competition for resource, and modelling life events discretely and probabilistically generates inter-individual variability, as well as geographical and demographic stochasticity, therefore accounting for several sources of uncertainty around population dynamics.

##### *Monitoring sub-model.*

We assumed that the manager makes no errors during monitoring, thus $N_t$ represents the exact population size at each time step. This assumption avoided modelled stochastic monitoring errors that would have challenged a full understanding of management dynamics.

##### *Decision-making sub-model.*

Manager and farmer decision-making is modelled in GMSE using evolutionary algorithms (Hamblin 2012). Each time an agent makes a decision, the GMSE evolutionary algorithm generates a set of random possible policies for managers (culling costs) or action plans for farmers (number of culls), and then allows this set to evolve on its own self-contained timescale. Policies or action plans that are better aligned to an agent's goal have a relatively high fitness, and the fittest ones are selected to be the agent's policy/action plan when the conditions for the algorithm termination are met (see supporting information S1 in Duthie et al. 2018, and GMSE documentation for further details). Our model thereby computes a practical but not necessarily optimal decision, recognizing that most people cannot think of every single possibility to choose the optimal one, but can choose the best option among those they could conceive. This process generates inter-individual variability, errors, and stochasticity in agents' decision-making, therefore simulating several sources of uncertainty around human behavior.

*Timing strategies implementation.*

CTL is the default strategy in GMSE: at each time step $t$, the evolutionary algorithm calculates an appropriate cost of culling (most likely a raise in the cost when $N_t < T_N$ and a decrease when $N_t > T_N$). In contrast, when applying ATI, the manager updates the policy only if $N_t$ is out of the permissive range ($T_N \pm P_T$). Hence, the evolutionary algorithm is called only if

$$\left|\frac{N_t}{T_N} - 1\right| > P_T$$

Otherwise, the cost is left the same as the previous time step. Lastly, when applying TRJ, the process is the same as ATI, except that the decision to update is based on a prediction of next time step's population size $\widehat{N}_{t+1}$ instead of $N_t$. We chose as a predicting function a simple linear extrapolation based on the current ($N_t$) and previous ($N_{t-1}$) population sizes that has the advantage of including the influence of the active policy on population variation in a simple way. Hence, with TRJ the condition for calling the evolutionary algorithm is

$$\left|\frac{\widehat{N}_{t+1}}{T_N} - 1\right| > P_T ,$$

With $\widehat{N}_{t+1} = N_t + (N_t - N_{t-1})$.

Otherwise, the cost stays the same as previous time step. After a time step without calling the evolutionary algorithm, the manager starts the next one with an addition of a proportion $B_b$ of $B_M$ b.u. to their regular budget $B_M$. (See Fig. A5 for a flowchart of the different strategies.)

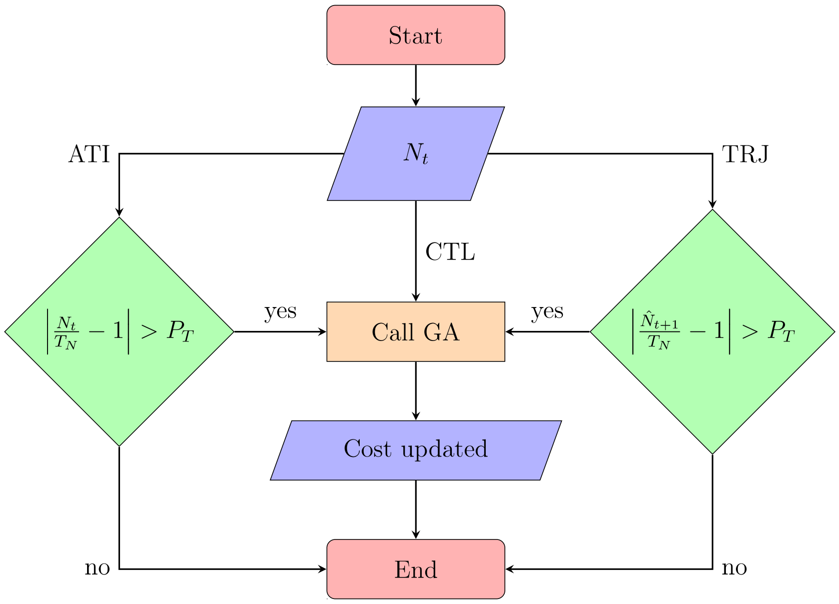


Fig. A4. Flowchart of the three timing strategies.

Table A4.1. Summary of useful symbols.

| Symbol | Status | Description | Unit |
|---|---|---|---|
| $t_{max}$ | constant | max simulation time | times steps |
| $T_N$ | constant | manager's target for population size | nb. of individuals |
| $N_0$ | constant | initial population size | nb. of individuals |
| $N_t$ | variable | population size monitored at time step t | nb. of individuals |
| $P_T$ | variable | permissiveness around $T_N$ | % of $T_N$ |
| $B_M$ | variable | manager's initial budget | b.u. |
| $B_b$ | variable | budget bonus amount | % of $B_M$ |
| $f_{ext}$ | outcome | extinction frequency over a set of replicates | % of replicates |
| $Y_{end}$ | outcome | average farmers' yield at the end of a simulation. | % of landscape max productivity |
| $Y_{ineq}$ | outcome | average differential between lowest and highest farmers' yields at the end of a simulation | % of highest yield |
| $d_T$ | outcome | Average distance between $Nt$ and $T_N$ at the end of a simulation | % of $T_N$ |
| $t_w$ | outcome | Average proportion of time steps without intervention | % of simulation time |

Table A4.2. GMSE parameter values. Parameters not mentioned here were set to default (as in https://confoobio.github.io/gmse/articles/SI3.html).

| Parameter | Value | Description |
|---|---|---|
| time_max | 20 | Maximum time steps in simulation |
| land_dim1 | 200 | Width of landscape (horizontal cells) |
| land_dim2 | 200 | Length of landscape |
| res_death_type | 0 | Rules affecting resource death (consumption-based) |
| res_birth_type | 0 | Rules affecting resource birth (consumption-based) |
| observe_type | 3 | Type of resource observation (transect observation) |
| res_move_obs | FALSE | Resource move during transect observation |
| res_consume | 0.5 | Pr. of a landscape cell's value reduced by the presence of a resource in a time step |
| max_ages | 5 | The maximum number of time steps a resource can persist before it is removed |
| minimumcost | 10 | The minimum cost of a farmer performing culling |
| user_budget | 1000 | A farmer's budget per time step for performing any number of actions |
| manager_budget | 1000 | A manager's budget per time step for setting policy |
| manage_target | 2000 | The manager's target resource abundance |
| RESOURCE_init | 1000 | The initial abundance of resources |
| culling | TRUE | Resource culling (removes a resource entirely) is a policy option |
| stakeholders | 40 | Number of farmers in the simulation |
| landownership | TRUE | farmers own land and increase utility indirectly from landscape instead of resource use |
| manager_sense | 0.15 | A metric of managers accuracy in predicting change in stakeholder behaviour given a change in cost |
| consume_surv | 4.75 | Amount of cell value for a resource to eventually survive until the next time step |
| consume_repr | 5 | Amount of cell value for a resource to eventually produce offspring |
| times_feeding | 12 | Maximum number of times a resource consumes landscape value per time step |
