## Supplementary material for "Intervene or wait? Modelling the timing of intervention in conservation conflicts adaptive management under uncertainty": French version of the abstract

### ABSTRACT (FR)

Le timing d'intervention des gestionnaires de biodiversité peut être déterminant dans le succès d'un programme de conservation, tout particulièrement quand leurs objectifs sont incompatibles avec des activités humaines (conflits de conservation). Mais l'incertitude associée aux systèmes socio-écologiques, ainsi que l'irréversibilité potentielle des conséquences d'un retard d'action peut pousser les gestionnaires à simplement intervenir dès que possible. Pourtant, y renoncer quand la situation le permet peut être bénéfique, notamment en mettant efficacement à profit les ressources non-utilisées. Nous proposons ici une stratégie basée sur le monitoring pour choisir si une intervention est nécessaire ou si attendre est préférable. Cette étude évalue la capacité de cette stratégie à satisfaire à la fois les objectifs de conservation et ceux des activités humaines en comparaison avec une stratégie d'intervention systématique et inconditionnelle. Pour ce faire, nous avons développé un modèle individu-centré de conflit de conservation entre des gestionnaires cherchant à conserver une population animale et des agriculteurs cherchant à en minimiser l'impact sur leurs cultures. Nous avons ensuite simulé une gestion adaptative du conflit sous contrainte budgétaire pour chaque stratégie, tout en prenant en compte l'incertitude associée à la dynamique de la population et à la prise de décision des parties prenantes. Quand la décision était basée sur une prédiction de la trajectoire de la taille de la population, notre stratégie était au moins aussi performante qu'une intervention inconditionnelle et permettait aux gestionnaires d'économiser des ressources en évitant des interventions non nécessaires. Lorsqu'un budget trop faible rendait la gestion difficile, notre stratégie a considérablement amélioré les résultats relatifs à la conservation en compensant le manque de ressources par les bénéfices accumulés au cours des périodes sans intervention. Ces résultats montrent que notre stratégie devrait être envisagée car elle peut assurer une gestion équitable du conflit tout en permettant une utilisation plus efficace des ressources de gestion, souvent limitantes en conservation de la biodiversité.

**Mots-clés**: Adaptive management; Conflits de conservation; Gestion de biodiversité; Management Strategy Evaluation; Modèle individu-centré; Modélisation de la prise de décision; Timing d’intervention; Incertitude.
